# PRG2 and AQPEP are misexpressed in fetal membranes in placenta previa and percreta

**DOI:** 10.1101/2020.08.14.248807

**Authors:** Elisa T. Zhang, Roberta L. Hannibal, Keyla M. Badillo Rivera, Janet H.T. Song, Kelly McGowan, Xiaowei Zhu, Gudrun Meinhardt, Martin Knöfler, Jürgen Pollheimer, Alexander E. Urban, Ann K. Folkins, Deirdre J. Lyell, Julie C. Baker

## Abstract

The obstetrical conditions placenta accreta spectrum (PAS) and placenta previa are a significant source of pregnancy-associated morbidity and mortality, yet the specific molecular and cellular underpinnings of these conditions are not known. In this study, we identified misregulated gene expression patterns in tissues from placenta previa and percreta (the most extreme form of PAS) compared with control cases. By comparing this gene set with existing placental single-cell and bulk RNA-Seq datasets, we show that the upregulated genes predominantly mark extravillous trophoblasts. We performed immunofluorescence on several candidate molecules and found that PRG2 and AQPEP protein levels are upregulated in both the fetal membranes and the placental disk in both conditions. While this increased AQPEP expression remains restricted to trophoblasts, PRG2 is mislocalized and is found throughout the fetal membranes. Using a larger patient cohort with a diverse set of gestationally aged-matched controls, we validated PRG2 as a marker for both previa and PAS and AQPEP as a marker for only previa in the fetal membranes membranes. Our findings suggest that the extraembryonic tissues surrounding the conceptus, including both the fetal membranes membranes and the placental disk, harbor a signature of previa and PAS that reflects increased trophoblast invasiveness.

**Summary sentence:** 3SEQ and immunofluorescence reveal that extravillous trophoblast factors, most notably PRG2 and AQPEP, define the diseases placenta previa and placenta accreta spectrum (PAS) in both the chorioamniotic membranes and the placental disk.

## Introduction

Placenta accreta spectrum (PAS) is a condition that is increasing at an alarming rate^1^, yet despite its significant harm to mothers and infants, the molecular and cellular origins are not well-understood. PAS is characterized by excessive invasion of the placental cells into the myometrium or muscle wall of the uterus. Deliveries associated with PAS are extremely challenging because the placenta becomes pathologically attached to the uterus, leading to hemorrhage and high rates of maternal and neonatal morbidity and mortality^1,2^. PAS is characterized by the subtypes accreta, increta, and and percreta delineating the extent of this invasion^1^. Percreta is the most extreme case, in which invasion extends through the uterine wall and sometimes into neighboring organs such as the bladder^1,3^. Nearly all accreta patients are also afflicted with previa, a condition in which the placenta is located in the lower segment of the uterus, directly covering the cervix^4,5^. This placement thus precludes vaginal delivery and puts the mother at risk of intra- and post-partum hemorrhage^6–9^. Both PAS and previa are associated with uterine damage or dysfunction as a result of previous cesarean deliveries, endometrial ablation, in vitro fertilization (IVF), or advanced maternal age^1,10–13^. A combination of multiple previous cesarean deliveries and previa elevates the risk of accreta to 67%, strongly supporting the correlation between uterine damage, previa, and PAS^5^.

While placental trophoblast cells appear to be at the heart of PAS, the role they play is unknown. The prevailing model is that uterine scar tissue, combined with placental placement in the lower uterus (previa), leads to deficient decidualization and the absence of appropriate regulatory signals to the trophoblasts, resulting in abnormal and aggressive placental invasion. Consistent with this model, PAS is characterized by expanded numbers of invasive extravillous trophoblasts (EVT)s^14–16^. EVTs are also expanded in previa, even though placental overinvasion is not characteristic of this disease^17^. The misexpression of several molecules within trophoblasts have been shown to correlate with disease, including lower levels of β-catenin in previa and accreta^18^, higher levels of HMG1 and VEGF in previa^19^, higher levels of DOCK4 in accreta trophoblasts^20^, and absence of osteopontin staining in percreta^21^. An in-depth examination of the involvement of trophoblasts, even years after uterine damage, is thus intriguing and critical for understanding disease etiology.

Although the placental disk plays a vital role in many diseases of pregnancy, the fetal membranes have also been implicated in some conditions. These extraplacental membranes, which line the fluid-filled amniotic sac to protect the fetus, are susceptible to infection and/or rupture that can jeopardize fetal health and lead to preterm birth. While not previously thought to be affected in placental diseases like preeclampsia, the fetal membranes have been found to contain cellular and molecular pathologies^22^. For example, in severe preeclampsia, the trophoblast layer is expanded in thickness in the fetal membranes. This is further accompanied by misexpression of many oxidative stress and immune genes, including key pregnancy proteins such as PAPPA, HTRA4, and CSH1^22^. In addition, recent studies report cases of physically adherent membranes during delivery, as well as the presence of myometrial myofibers attached to these membranes^15,23^. These cases are often associated with either concurrent, subsequent, or prior PAS, thus raising the idea of fetal membrane involvement in this disease^24^. Therefore, while trophoblast overinvasion and expansion of EVTs in PAS is seen primarily in the placental disk^16^, the membranes may also be compromised.

Here we examined the fetal membranes of PAS and previa compared to gestationally age-matched controls to identify molecules that could discriminate between these conditions. We found that 74 transcripts were significantly upregulated in cases of percreta and previa. These genes are highly enriched for extracellular molecules that are EVT-specific markers. We validated two of these molecules, Proteoglycan 2 (PRG2) and Aminopeptidase Q (AQPEP; otherwise known as Laeverin), using immunofluorescence in the membranes of PAS, previa, and a diverse set of controls. We further demonstrate that PRG2 and AQPEP are misexpressed within the placental disc in PAS. Overall, there is an EVT molecular signature within previa and PAS membranes, revealing that extraembryonic tissues, including the fetal membranes, are affected in these diseases.

## Methods

### Patient samples

All sample collection and clinical data abstraction was conducted in accordance with protocols approved by the Stanford Institutional Review Board (IRB #20237). Samples were processed and stored as formalin-fixed, paraffin embedded (FFPE) tissue blocks by the Stanford Pathology department. All samples included in this study and corresponding clinical data are listed in Supplemental Table S1. Two cohorts of patients were included in this study: the initial discovery cohort (cohort 1) of placenta previa, PAS, and spontaneous preterm birth (SPTB) control samples on which we performed 3SEQ and immunofluorescence (see below), and a validation cohort (cohort 2) composed of placenta previa, PAS, and gestationally age-matched controls (see below and see Results).

### 3SEQ and differential gene expression analysis in cohort 1 fetal membranes

Tissue regions containing all layers of the fetal membranes (amnion, chorionic mesoderm, trophoblasts, and decidua) were cored from FFPE blocks. RNA was isolated from these cores using the RecoverAll Total Nucleic Acid Isolation Kit (Ambion, cat. no. 1975). 3SEQ libraries were prepared according to Beck et al. 2010^25^. Briefly, polyA selection was performed using the Dynabeads mRNA Purification Kit (Invitrogen), followed by cDNA synthesis using a P7-containing primer and Superscript III Reverse Transcriptase (Invitrogen) and A-tailing. After purification using the MinElute Kit (Qiagen) and ligation of double-stranded P5 linker to cDNA, agarose gel fractionation and purification (MinElute Kit; Qiagen) was performed to isolate linker-ligated cDNA of fragment length 200-300 bp. Final libraries were generated by PCR amplification of the linker-ligated cDNA with Illumina-compatible primers and Phusion PCR Master Mix (New England Biolab) using a 15 cycle program (98°C for 30 sec; 15 cycles of 98°C for 10 sec, 65°C for 30 sec, 72°C for 30 sec; 72°C for 5 min). 3SEQ libraries were sequenced using an Illumina HiSeq 2500 to an average read depth of ∼25 million reads per sample. Sequencing read statistics are provided in Supplemental Table S2. Single-end reads were mapped to the hg38 reference genome (Supplemental Table S2) using TopHat^26^. Adapters were removed using cutadapt^27^, and read quality control was performed using FastQC. Gene expression counts were obtained using UniPeak^28^. The sequencing reads identified a total of 11,476 genes. Of these transcripts, we excluded 168 genes with fewer than an average of 10 reads per sample across all samples. Of the remaining 11,308 transcripts, we excluded 329 instances of ambiguous multi-gene mappings, leaving a total of 10,979 genes for subsequent analysis. Differential gene expression was analyzed using DESeq^29^. GO term analysis was performed using geneontology.org with Fisher’s exact test and Bonferonni correction. 3SEQ data from this study is available via the NCBI dbGaP under accession no. phs002075.v1.p1. Genes differentially-expressed between SPTB and both previa and PAS were compared against genes differentially-expressed between MACS-isolated HLA-G^+^ extravillous trophoblasts (EVTs) and EGFR^+^ villous cytotrophoblasts (VCTBs) from first trimester placentas from 10-12 weeks of gestation, as described in detail in Vondra et al. 2019^30^. RNA-seq data from Vondra et al. 2019 was obtained from GEO Series accession number GSE126530.

### Histology and Immunofluorescence on cohorts 1 and 2

10% formalin-fixed, paraffin-embedded samples of fetal membranes and placental disks from cases and controls of cohorts 1 and 2 from the Stanford Pathology archive were cut into 5 μm-thick sections. Sections were stained with hematoxylin and eosin (H&E) by deparaffinization with xylenes, rehydration in a graded series of ethanol in distilled water, staining with Harris hematoxylin (Sigma-Aldrich cat. no. SLBV6928), dehydration in a graded series of ethanol to 70%, staining with alcoholic eosin Y (Sigma-Aldrich cat no. SLBW3770), followed by further dehydration and subsequent mounting in Permount (Fisher Scientific cat no. SF15-100). H&E staining was performed to identify regions with similar cell type compositions (amnion, chorionic mesoderm, trophoblasts, and decidua), which were then cored and subsequently subjected to either 3SEQ (see above) or re-embedding in paraffin for immunofluorescence.

Immunofluorescence was performed on cut sections after antigen retrieval in 0.01 M sodium citrate buffer, pH 6 in a pressure cooker. Sections were then blocked with blocking reagent (5% goat serum, 1% BSA, and 0.1% Triton-X 100 in PBS) and stained overnight at 4° C with antibodies against PRG2 (1:1000 dilution, rabbit; Sigma-Aldrich cat. no. HPA038515), HLA-G (1:10,000 dilution, mouse; Santa Cruz cat. no. sc-21799), or AQPEP (1:1000 for single-marker staining, 1:2000 for co-staining with anti-CK7; rabbit, Abcam cat. no. ab185345). Immunofluorescence experiments were performed semi-quantitatively by titrating the primary antibody concentration to a point at which the which the lowest levels of the protein remained detectable, thus ensuring that the dynamic range of tissue intensities was well-represented. After successive washes in PBST (0.1% Tween-20 in phosphate-buffered saline), native peroxidase activity was quenched with 3% hydrogen peroxide in PBS for 30 minutes. Biotin-conjugated secondaries against the relevant species (1:5000 goat anti-rabbit; 1:1000 goat anti-mouse) were applied for 1 hr. at room temperature (Jackson Immunoresearch, cat. no. 111-065-144 and 111-065-166, respectively). All antibodies were diluted in blocking reagent. After successive washes to remove unbound secondary antibody, an ABC peroxidase staining kit (Thermo Scientific Pierce, cat. no. 32020) and tyramide signal amplification (TSA) kit were used to apply Cy3 fluorophore (Perkin Elmer cat. no. NEL744001KT) for single-marker staining.

Where indicated, double-marker staining was performed sequentially after staining with the marker of interest, followed by repeating all steps from antigen retrieval through TSA using an antibody against CK7 (1:200, mouse; Dako cat. no. MS-1352) and the FITC fluorophore (Perkin Elmer cat. no. NEL741001KT). Slides were mounted in ProLong Gold Antifade Mountant with DAPI (Life Technologies, P36931). Images were acquired on a Leica DMRXA2 inverted fluorescence microscope and a Nikon Eclipse Ti inverted confocal microscope. For HLA-G, an additional 2 SPTB samples, as well as 1 increta sample that was not included in the original 3SEQ, were included amongst cohort 1 immunofluorescence analyses.

### Immunofluorescence image analysis

Image intensity quantification and normalization were performed using ImageJ^32^. For discovery cohort samples (Fig. 2), tissue regions were selected by performing Otsu thresholding in ImageJ on the DAPI channel, from which a mask was created to quantify mean fluorescence intensity of this entire masked tissue region in the marker (PRG2, AQPEP, or HLA-G) channel. For the decidua basalis samples (Fig. 3), additional analyses were performed using Otsu thresholding on the channel of the marker to identify regions with marker staining, from which a mask was created to quantify mean fluorescence signal intensity specifically from regions positive for marker staining. For cohort 2 samples (Fig. 4), regions of interest (ROIs) containing CK7+ trophoblasts were selected manually, and total fluorescence intensity for PRG2, AQPEP, or HLA-G was calculated for each trophoblast-containing ROI. HLA-G^+^ / CK7^+^ cell counting data were obtained using custom MATLAB scripts.

**Figure 1:**
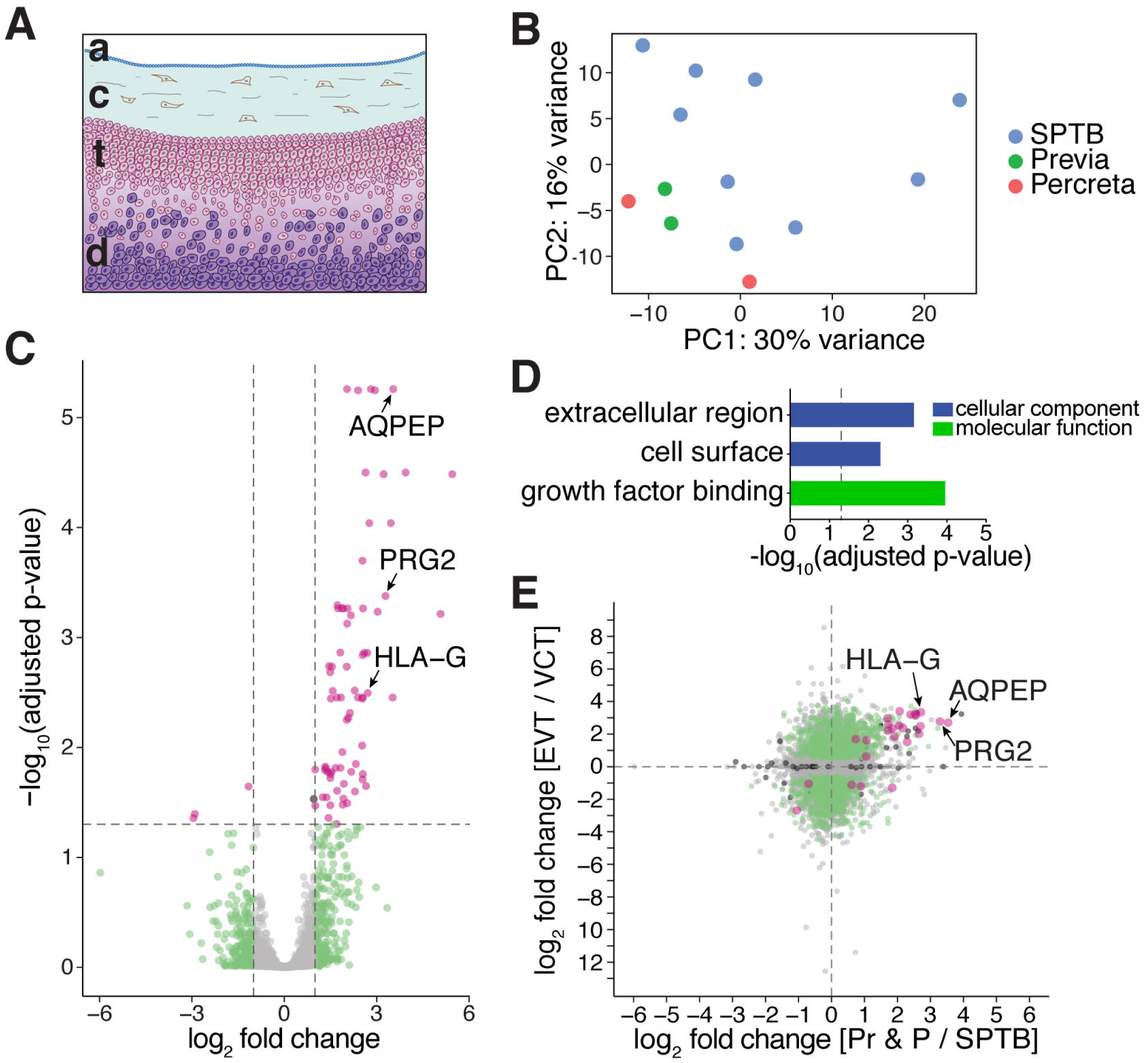
Genes upregulated in previa and percreta are EVT-specific. A. Structure and cell types of human fetal membranes. Schematic illustration showing the cellular layers that comprise the chorionic membranes: the amnion (a), chorionic mesoderm (c), trophoblasts (t), and decidua (d). B. Principal component analysis of 3SEQ of all 13 samples: 2 previa, 2 percreta, and 9 spontaneous pre-term birth (SPTB). C. Volcano plot showing differentially-expressed genes (DEGs) between previa and percreta samples vs. SPTB samples. Genes demonstrating both an adjusted p-value < 0.05 and a log_2_ fold-change either < −1 or > 1 are indicated in magenta. Genes with only a log_2_ fold-change either < −1 or > 1 are indicated in green, whereas genes with only an adjusted p-value < 0.05 are indicated in black. All remaining genes that reached neither statistical significance nor fold-change thresholds are indicated in gray. D. GO terms associated with previa and percreta vs. SPTB DEGs. −log_10_ Bonferonni-corrected adjusted p-values are shown. Dashed gray line indicates adjusted p-value < 0.05 threshold. E. Scatter plot illustrating the overrepresentation of invasive extravillous trophoblast (EVT) genes amongst genes upregulated in previa and percreta (Pr & P) vs. SPTB (magenta). Genes only significantly up- or down-regulated in previa & percreta vs. SPTB but not in EVTs vs. progenitor villous cytotrophoblasts (VCTs) are indicated in black. Genes only significantly up- or down-regulated in EVTs vs. VCTs but not in previa & percreta vs. SPTB are indicated in green. All other genes are indicated in gray.

**Figure 2:**
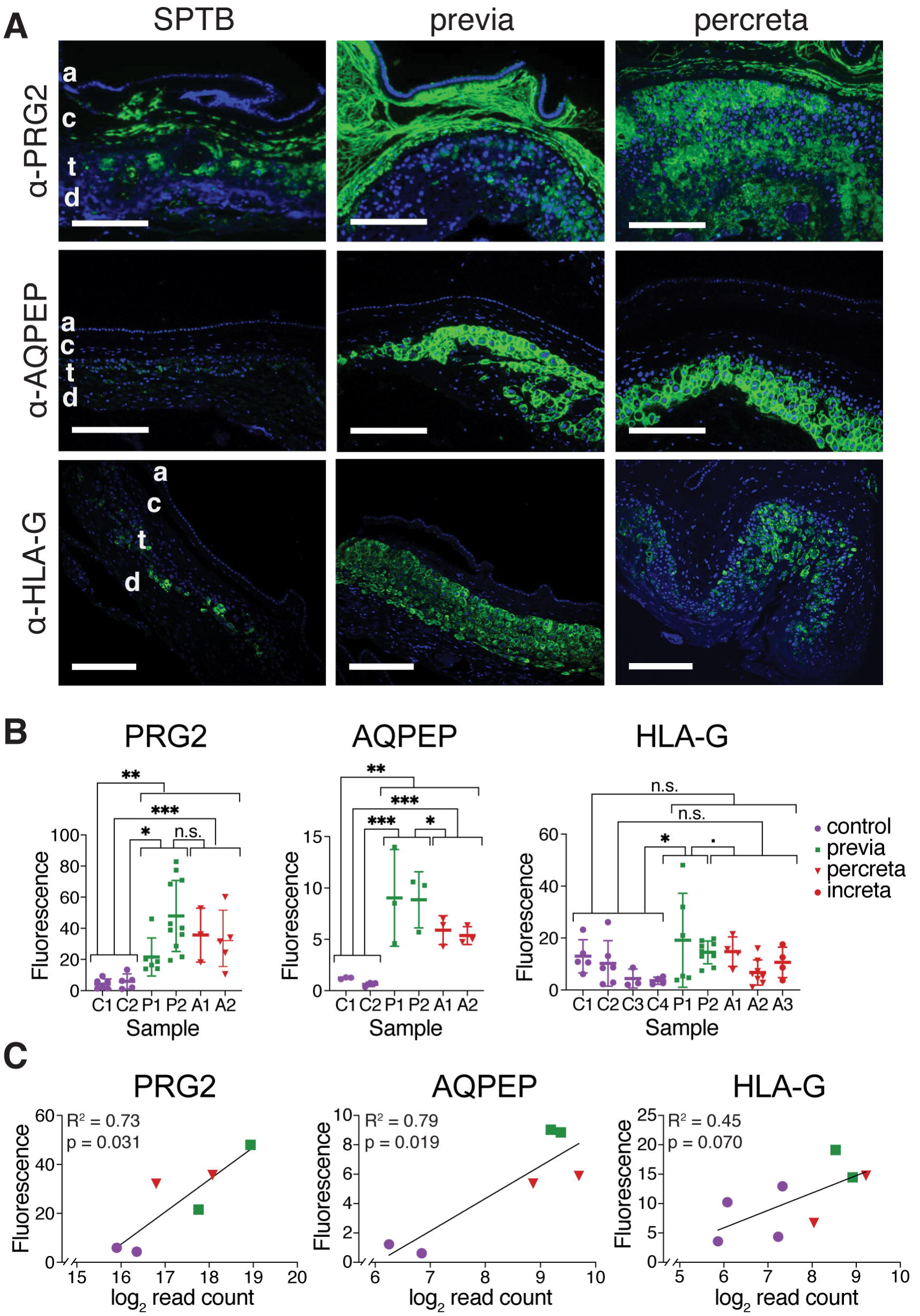
Higher levels of PRG2 and AQPEP protein in previa and PAS fetal membranes. A. Representative images from immunofluorescence staining of SPTB, previa, and percreta chorionic membrane samples using anti-PRG2, anti-AQPEP, and anti-HLA-G antibodies. Scale bar = 200 μm. blue = DAPI, green = antibody staining. B. Quantification of the mean fluorescence intensities (Fluorescence) per image (see Methods) for anti-PRG2 (left), anti-AQPEP (middle), and anti-HLA-G (right). p-value < 0.05 (*), p-value < 0.01 (**), p-value < 0.001 (***), p-value < 0.1 (.), and not statistically significant (n.s.) are indicated as shown. C. Linear regression of 3SEQ RNA levels (log_2_ normalized read count) vs. average sample protein levels as measured by immunofluorescence intensity (Fluorescence) for the indicated markers PRG2, AQPEP, and HLA-G.

**Figure 3:**
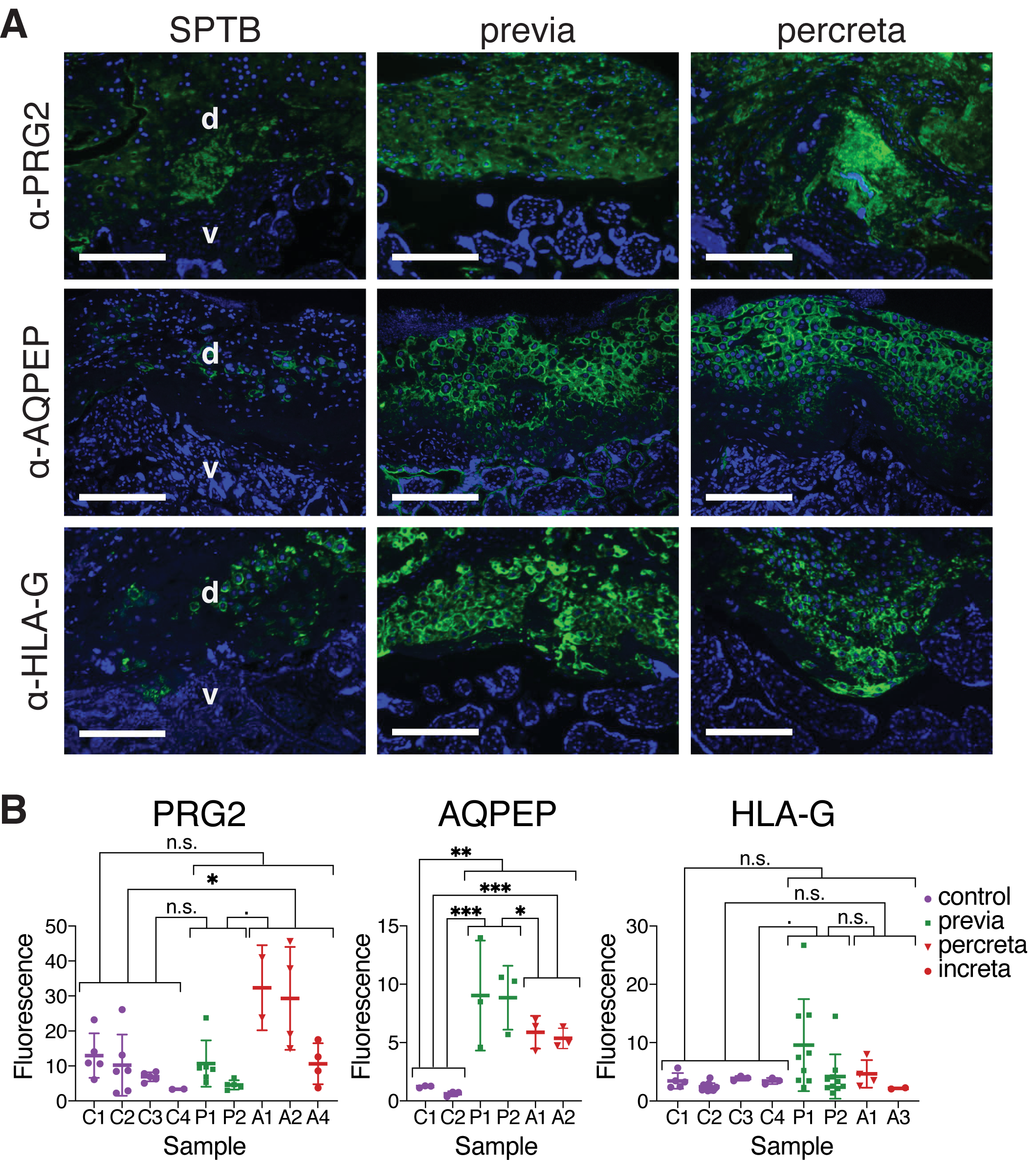
PRG2 and AQPEP protein levels are higher in previa and PAS placental disk samples compared to SPTB. A. Representative images from immunofluorescence staining of SPTB, previa, and percreta placental disk samples for the markers PRG2, AQPEP, and HLA-G. Scale bar = 200 μm. blue = DAPI, green = antibody staining. Villi (v) and decidua (d) are indicated. B. Quantification of anti-PRG2, anti-AQPEP, and anti-HLA-G mean fluorescence intensity levels (Fluorescence) per image (see Methods). p-value < 0.05 (*), p-value < 0.01 (**), p-value < 0.001 (***),p-value < 0.1 (.), and not statistically significant (n.s.) are indicated as shown.

**Figure 4:**
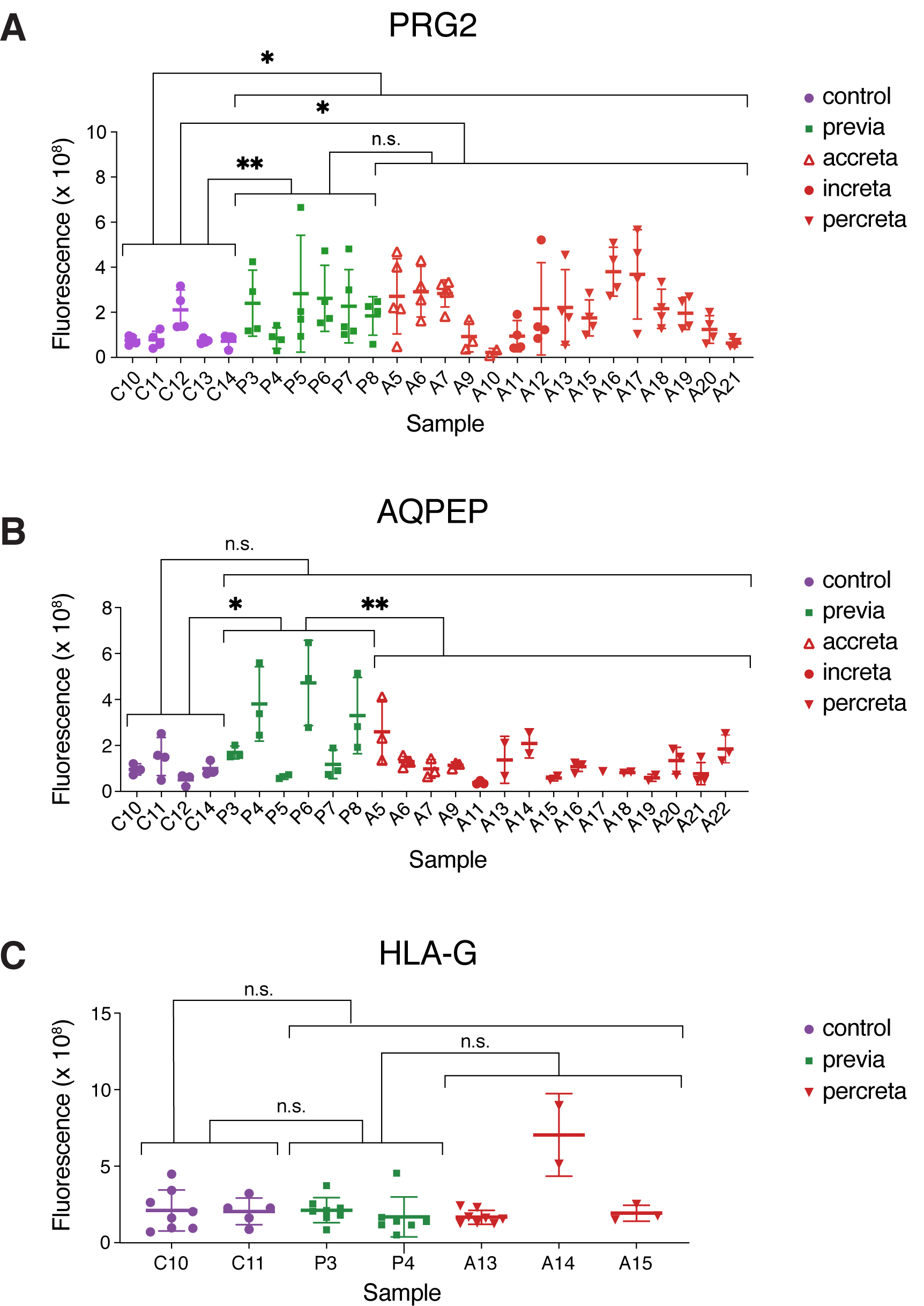
Validation cohort of patients display higher levels of PRG2 in both previa and accreta and higher levels of AQPEP in previa. A. Quantitation of PRG2 total immunofluorescence signal intensities in CK7^+^ trophoblasts. B. Quantitation of AQPEP total immunofluorescence signal intensities in CK7^+^ trophoblasts. C. Quantitation of HLA-G total immunofluorescence signal intensities in CK7^+^ trophoblasts. (A-C) p-value < 0.05 (*), p-value < 0.01 (**), p-value < 0.001 (***), and not statistically significant (n.s.) are indicated as shown.

Since protein marker levels in each patient sample was measured using multiple images, linear mixed-effects regression analysis was performed in R to assess statistical significance of fluorescence intensity differences between patient groups, with the disease state as the fixed effect and the patient donor as the random effect. Specifically, we used ANOVA to test the model of fixed effects (disease) and random effects (patient) fluorescence ∼ disease + (1|patient) against the null model fluorescence ∼ (1|patient).

Linear mixed-effects regression analysis was performed in R to assess the correlation between mRNA transcript levels as measured by 3SEQ and marker protein levels as determined by immunofluorescence. We used ANOVA to test the model fluorescence ∼ transcript_level + (1|patient) against the null model fluorescence ∼ (1|patient).

## Results

### Molecular differences in fetal membranes between previa, percreta, and spontaneous preterm birth

To identify molecular signatures of previa and percreta in the fetal membranes, we performed 3SEQ on biobanked paraffin tissue blocks from previa (2), percreta (2) and SPTB (9) samples at 28-32 weeks of gestation (see Methods)^25^. We first used histopathology to identify tissue regions that contained similar cellular compositions and tissue architecture (amnion, chorionic mesoderm, trophoblasts, and decidua; see Fig. 1A for schematic). We then cored these regions from these tissue blocks for RNA extraction and sequencing. Principal component analysis (PCA) using data for all 13 samples show that these samples do not cluster by disease state (Fig. 1B), suggesting that these conditions are not globally distinct from one another. To identify individual genes that distinguish previa and percreta from SPTB, we performed differential gene expression analysis, which revealed 77 misexpressed genes, with 74 upregulated in previa and percreta relative to SPTB and 3 downregulated in previa and percreta (Fig. 1C). Overall, while the fetal membranes are not globally different between these conditions, there are select genes that distinguish previa and percreta from SPTB, suggesting that these might be informative biomarkers.

### Genes misexpressed in previa and percreta are cell surface molecules expressed by extravillous trophoblast factors

To examine whether the 74 upregulated genes can provide insight into the biology of these diseases, we performed GO analysis (Fig. 1D). Despite the low number of genes in this dataset, we found significant enrichment for extracellular, cell surface proteins and embryonic growth factors. Of the 71 upregulated genes that had GO annotations, 55% (39/71) are extracellular. These extracellular annotations include the terms extracellular region (34/71; p=.00066), plasma membrane (14/71; p=.0047), and growth factor binding (8/71; p=0.00011). Molecules represented include the HLA class I histocompatibility antigen, alpha chain G (HLA-G), proteoglycan 2 (PRG2), lysophosphatidic acid receptor 2 (LPAR2), ephrin-B1 (EFNB1), allograft inflammatory factor 1-like protein (AIF1L), fibroblast growth factor receptor 1 (FGFR1), the VEGF receptor FLT1, IL-2 receptor beta (IL2RB), and the leukemia inhibitory factor receptor (LIFR). Therefore, we show that genes aberrantly upregulated in previa and percreta relative to SPTB are predominantly extracellular proteins, some of which are involved in growth factor signaling.

To determine which placental cell types are involved in previa and PAS, we compared the 74 upregulated genes to markers of specific cells found in the placenta. We first used a recent single-cell study that identified 194 genes that could discriminate between different placental cell types, including EVTs, syncytiotrophoblasts, decidual cells, endothelial cells, and immune cells^33^. When we compared the 74 upregulated genes to the 194 specific cell-type markers, we identified 9 genes that are present in both studies. Of these 9 genes, 8 (88.9%) are specific to EVTs (Supplemental Table S3). Since only 11.9% (23 out of 194) of the cell type markers overall are specific for EVTs, this comparison demonstrates that genes upregulated in previa and percreta are highly enriched (∼89%) for EVT-specific markers (test of two proportions, p-value = 6.4 × 10^−9^).

To further support that the upregulated signature represents EVTs, we examined an RNA-seq^30^ and a microarray dataset^34^, both of which identified transcripts specific to EVTs relative to their progenitor cells, the villous cytotrophoblasts (VCTs). We first overlapped the 74 genes upregulated in previa and percreta with both datasets. We found 42 and 21 of the 74 genes in the RNA-seq and microarray datasets, respectively. Comparison with EVT- and VCT-specific genes from the RNA-seq experiment showed that 95.2% (40/42) were enriched in EVTs while only 4.8% (2/42) were enriched in VCTs (40/42 EVT genes vs. 1159 EVT genes / 1990 total EVT-VCT genes, test of two proportions p = 3.1 × 10^−6^; Fig. 1E, Supplemental Table S3). Comparison with EVT- and VCT-specific transcripts from the microarray experiment showed that 95.2% (20/21) were upregulated in EVTs versus only 4.8% (1/74) upregulated in VCTs (20/21 EVT genes vs. 428 EVT genes / 885 total genes, test of two proportions p = 5.7 × 10^−5^; Supplemental Table S3). In aggregate across all 3 datasets, including single-cell RNA-seq, RNA-seq, and microarray, 91.8% of the overlapping transcripts (45/49) are EVT-enriched whereas only 8.2% (4/49) are enriched in non-EVTs (Supplemental Table S3). Overall, the genes in the fetal membranes that distinguish previa and percreta from SPTB have a strong EVT transcriptional signature.

### PRG2 and AQPEP are more abundant in previa and percreta compared to SPTB

We next examined the protein expression levels and localization of several of the upregulated genes in histological sections of fetal membranes from the previa, percreta, and SPTB patient samples profiled above. We performed immunofluorescence using antibodies against AQPEP (aminopeptidase Q or Laeverin), PRG2 (proteoglycan 2), and HLA-G (human leukocyte antigen G). In order to examine changes in protein levels as quantitatively as possible, we used the minimum antibody concentration at which the lowest levels of protein was just barely detected, thus ensuring that the highest levels of each protein did not exceed saturation and remained in linear range (see Methods). Consistent with the 3SEQ data, we observed an increase in staining of these antibodies in previa and percreta samples compared to SPTB (Fig. 2A). We found that whereas PRG2 is localized to both the trophoblasts and the chorionic mesoderm in SPTB, it is widely expressed throughout the membranes in all layers except the amnion in previa and percreta (Fig. 2A, top panel). In contrast, AQPEP and HLA-G are localized specifically to EVTs in all samples - SPTB, previa, and percreta. However, the intensity of staining within the EVTs is increased in the previa and percreta samples (Fig. 2A, middle and bottom panels).

To quantify differences in AQPEP, PRG2 and HLA-G protein levels between previa, percreta, and SPTB, we measured mean fluorescence intensities of 3-9 tissue regions imaged from each of 6-9 patients and performed linear mixed-effects regression. We found that PRG2 and AQPEP are overexpressed in both previa and percreta compared to SPTB (p = 0.0051 and p = 0.0012, respectively; Fig. 2B). In addition, AQPEP expression is higher in previa than percreta (p = 0.027), and HLA-G expression is higher in previa compared to SPTB (p =0.023) but does not reach statistical significance when compared to PAS (p = 0.059) (Fig. 2B). Co-staining with anti-HLA-G and with anti-cytokeratin-7 (CK7) to mark all trophoblasts (Supplemental Fig. S1A) revealed that fetal membrane samples from previa patients contained more HLA-G^+^ trophoblasts compared to percreta, increta, or SPTB. This suggests that in these previa samples, the higher levels of HLA-G reflect an excess of trophoblasts (Supplemental Fig. S1B). Taken together, this cohort analysis suggests that PRG2 and AQPEP are elevated in both previa and percreta, with PRG2 also being mislocalized throughout the fetal membranes (Fig. 2, Table 1).

**Table 1.**

|  | Cohort #1 M (v. SPTB) |  | Cohort #2 M (v. ctrls) |  | Cohort #1 Plac (v. SPTB) |  |
| --- | --- | --- | --- | --- | --- | --- |
|  | Previa | PAS | Previa | PAS | Previa | PAS |
| PRG2 | + | + | + | + | - | + |
| AQPEP | + | + | + | - | + | + |
| HLA-G | + | - | - | - | - | - |

Next, we tested whether there was a direct correlation between RNA and protein levels for PRG2, AQPEP, and HLA-G. To this end, we plotted normalized read counts from the 3SEQ data against the average immunofluorescence staining intensity from the same patient and performed linear mixed-effects regression. The increases in protein and mRNA levels for PRG2 and AQPEP were strongly correlated, with an R^2^ value of 0.73 (p-value = 0.031) for PRG2 and R^2^ = 0.79 (p-value = 0.019) for AQPEP (Fig. 2C). HLA-G protein and mRNA levels were moderately correlated, though this did not reach statistical significance (R^2^ = 0.45, p-value = 0.069; Fig. 2C). Prior reports in the literature find that cellular concentrations of many proteins correlate only moderately with transcript levels^35,36^, with one review citing R^2^ values around 0.4^37^. Therefore, the strong correlations between the protein and RNA levels of these three genes support the idea that the remaining misregulated genes are also likely misregulated as proteins within the fetal membranes.

### PRG2 and AQPEP are more abundant in the placental disk in previa and PAS

We next asked whether the misregulation of PRG2, AQPEP, and HLA-G in the fetal membranes is also observed in the placental disk. To this end, we performed immunohistochemistry on the placental disk from the same pregnancies transcriptionally profiled by 3SEQ (Fig. 3). In all comparisons, AQPEP is significantly more abundant in previa and percreta than in SPTB, consistent with what was described for the fetal membranes (Fig. 3A,B). Protein levels for HLA-G were elevated in previa vs. SPTB, matching the trend observed in the fetal membranes, though without passing a significance threshold (p=0.059; Fig. 3B). In contrast to AQPEP and HLA-G, which display the same overall trends between the placental disk and the membranes, PRG2 protein levels can discriminate between SPTB and percreta, but not between SPTB and previa (Fig. 3A,B). PRG2 levels are also higher in percreta than in previa, though without statistical significance (p=0.079). We thus find that many of the molecular changes in the fetal membranes between disease states are also present in the placental disk.

### Second patient cohort supports findings for PRG2 and AQPEP

To determine whether PRG2, AQPEP, and HLAG could be used more broadly to discriminate disease states, we performed immunofluorescence on a second, larger cohort of patient samples. This validation cohort (cohort 2) was composed of 5 gestationally age-matched control samples (31-37 wk), 6 previa samples (29-38 wk), and 16 PAS samples (4 accretas, 31-35 wk; 2 incretas, 31-36 wk; and 10 percretas, 28-37 wk). Importantly, the controls were from a variety of conditions that were not SPTB: cervical cancer requiring a C-section hysterectomy (n=2, 34 wk), term pregnancy with chorioamnionitis (n=1, 37 wk), and twin pregnancies in which one twin was affected with increta (n=1, 31 wk) or accreta (n=1, 32 wk; Fig. 4, Supplemental Fig. S2, Supplemental Table S1). In this larger cohort, we performed immunofluorescence by costaining with anti-CK7 to mark all trophoblasts and either anti-PRG2, anti-AQPEP, or anti-HLA-G (Supplemental Fig. S2). We then quantified the intensity of each marker in regions containing CK7^+^ trophoblasts as described in the Methods. In concordance with the results from the initial discovery cohort (Fig. 2A,B), protein levels of PRG2 are significantly higher in previa and PAS in all of the following comparisons: both previa and PAS compared to controls (p-value = 0.023), previa compared to controls (p=0.0099), and PAS compared to controls (p=0.038) (Fig. 4A). For AQPEP, we observe higher levels in previa relative to either controls (p = 0.050) or PAS (p = 0.0038), thus recapitulating the results of cohort 1 (Fig. 4B, Fig. 2A,B). However, in contrast to cohort 1, AQPEP levels in cohort 2 are not higher in PAS compared to controls. HLA-G did not recapitulate the higher levels or greater cell numbers in previa observed in cohort 1 (Fig. 4C, Fig. 2A,B). Overall, this second cohort, using different controls, validate all of the initial findings for PRG2 as well as a key trend for AQPEP (Fig. 2, Fig. 4, Table 1).

## Discussion

In this study, we identified several candidate molecules that are misexpressed in the fetal membranes of previa and PAS patients. These differences were initially identified by sequencing of patient samples and then confirmed by immunofluorescence using both discovery and validation cohorts. We found that PRG2 is upregulated in both previa and PAS patients in both cohorts, making this the most robust marker to emerge from this study. We further demonstrated that AQPEP discriminates between controls and previa in both cohorts. AQPEP also discriminates between controls and PAS in the discovery cohort but not in the second cohort. Data suggest that HLAG may be misexpressed in previa fetal membranes compared to controls, but this association is weaker than for PRG2 and AQPEP. The few differences in the observations between discovery cohort 1 and validation cohort 2 (HLA-G levels in previa, AQPEP levels in PAS) may be attributed to the inherent variability in human patient samples as well as the larger span of gestational ages in cohort 2 (see Results and Supplemental Table S1). The trends identified in the fetal membranes were also shown to exist in the placental disk of the discovery cohort. Taken together, this work suggests that PRG2 and AQPEP are markers for previa and PAS disease within the fetal membranes and further supports the idea that extraembryonic tissues are globally altered within these disease states.

PRG2 and AQPEP have previously been shown to be expressed in the placenta and have been implicated in diseases affecting pregnancy. PRG2 (Proteoglycan 2) is expressed in multiple cell types, with its highest expression in the placenta^38^. While the precise role of PRG2 during pregnancy is unknown, PRG2 has been shown to be higher in expression in preeclampsia^39–41^, maternal obesity^42^, gestational hypertension^43^, and Chagas disease^44^. PRG2 in its proform is also found at high levels in the placenta^38^ and in pregnancy serum^45^, where it forms a complex with PAPPA (pregnancy-associated plasma protein A), a molecule that has been implicated in placental development and pregnancy disorders such as low birth weight^46^, miscarriage^47^, and preeclampsia^48^. AQPEP (aminopeptidase Q or laeverin) is a placenta-specific aminopeptidase expressed in HLA-G^+^ EVTs and has been shown to be necessary and sufficient for EVT migration in in vitro primary villous explant outgrowth assays^49,50^. In addition, AQPEP inhibits the activity of other placental proteins by cleaving the N-terminal amino acids of peptides such as angiotensin III, kisspeptin-10, and endokinin C, suggesting that the enzyme plays important roles in human placentation^51^. AQPEP expression is higher in preeclampsia^52,53^, suggesting its involvement in placental disorders that warrants further investigation. Our identification of PRG2 and AQPEP as a feature of these diseases provides an inroad to more mechanistic studies.

The set of genes upregulated in previa and PAS are enriched for EVT-specific markers, suggesting that there may be a cell fate change toward a more invasive trophoblast program in these diseases. The presence of these molecular changes in previa and PAS tissues is hardly surprising, given that abnormal placentation into the poorly vascularized lower uterine segment may induce compensatory responses that are ultimately manifested at the structural, cellular, and molecular levels^54^. Whether these EVT changes are merely the consequence of an altered uterine environment and play no role in disease etiology, or whether they arise from an altered uterine environment and directly contribute to disease progression, remain to be determined. Defining the precise etiologies of these diseases will be invaluable for understanding disease progression and identifying possible points of intervention.

This work uncovers a disease signature in the fetal membranes of previa and PAS patients, an unexpected finding since the disease pathology for previa and PAS is manifested in the placental disk. This result is bolstered by clinical observations that support the possibililty of a fetal membrane component in these conditions. For example, some reports have noted adherent or retained membranes associated with PAS deliveries^23^. In some pregnancies, myofibers can be found associated with fetal membranes in conjunction with basal plate myofibers^24^, which are often associated with or precede PAS pregnancies^55,56^. Indeed, consistent with these reports, clinical notes from two of the PAS patients in our study (samples A1 and A15) indicated membranes adherent to the uterine walls. This raises the question of how the fetal membranes could harbor an invasive or “sticky” predisposition when the diseases have previously only been associated with the placenta itself. Three models might explain this phenomenon: 1) early implantation in the lower uterine segment (previa) results in a molecular perturbation in all extraembryonic tissues throughout the conceptus that is maintained for the duration of the pregnancy, 2) direct molecular communication between the fetal membranes and placental disk throughout gestation allow pathollogies in any one region to spread throughout the extraembryonic tissues, and 3) perturbed endocrine signaling due to previa and/or PAS that results in shared molecular signatures across the extraembryonic tissues. Recently, molecular and cellular changes were also found in preeclamptic fetal membranes^31^, which might support wider communication between different extraembryonic tissues either at implantation or throughout pregnancy.

Overall, this study is the first to examine molecular changes in the fetal membranes in previa and PAS and supports further study of the role of the fetal membranes in human disease. Future studies taking a comprehensive view of dysregulated molecules within the conceptus both at the site of pathological adherence as well as throughout the entire extraembryonic tissues, would be important for advancing our understanding of these diseases.

## Acknowledgements

We thank Amy Heerema-McKenney for assistance with identifying patients for this study and selecting tissue regions for 3SEQ. We thank Jessica K. Chang for assistance with bioinformatic analyses, Huaying Fang and Sunny X. Tang for advice on statistical analyses, and Kristen L. Wells for assistance with R graphics. We thank Opher Kornfeld for the MATLAB cell counting script. We thank Amarjeet Grewall for assistance with sectioning and histology. We thank Imee Datoc and Emily E. Ryan for assistance with patient chart data abstraction. We also thank Kelly Ormond for input on patient confidentiality-related matters. Lastly, we wish to thank all members of the Baker lab for suggestions and feedback regarding the project.

## Conflict of interest

The authors have no conflicts of interest to declare.

## Author Contributions

E.T.Z., J.R.S., and R.L.H. performed bioinformatic experiments and analyses. A.K.F. and D.J.L. identified patients for this study, biobanked patient samples, and provided expertise and guidance on study design. A.K.F. gathered the cases and controls and reviewed H&E slides to determine tissue areas to be cored for the project. R.L.H. performed 3SEQ. E.T.Z., K.M.B.R., and K.M. performed immunofluorescence experiments with assistance from X.Z. with data analysis. G.M., X.Z., J.P., M.K., and A.E.U provided EVT and VCT data prior to publication. G.M., X.Z., R.L.H., J.P., M.K., and E.T.Z. performed EVT and VCT data analysis and interpretation. J.P., and M.K. provided expertise on EVT biology. E.T.Z. and J.C.B. wrote the paper. All authors contributed to experimental design and interpretation and provided comments on the manuscript.

## Notes

Grant support: This work was funded by an NIH grant (NICHD R01 HD094513), a Stanford Discovery and Innovation Foundation Grant, and a Stanford SPARK grant to J.C.B. E.T.Z was supported by an A.P. Giannini Foundation postdoctoral fellowship, a Stanford Child Health Research Institute postdoctoral award, and a Stanford Dean’s Postdoctoral Fellowship. J.H.T.S. was supported by a National Science Foundation Graduate Research Fellowship and Stanford Graduate Fellowship. M.K. and J.P. are supported by grants from the Austrian Science Fund (P31470-B30 and P-33485-B30). D.J.L. was supported by The Arline and Pete Harman Faculty Scholar Fund from the Stanford Maternal Child Health Research Institute and the former Stanford Child Health Research Institute.

### Competing Interest Statement

The authors have declared no competing interest.

