## Supplementary figures and images for "PRG2 and AQPEP are misexpressed in fetal membranes in placenta previa and percreta"

### Supplemental Fig. S1

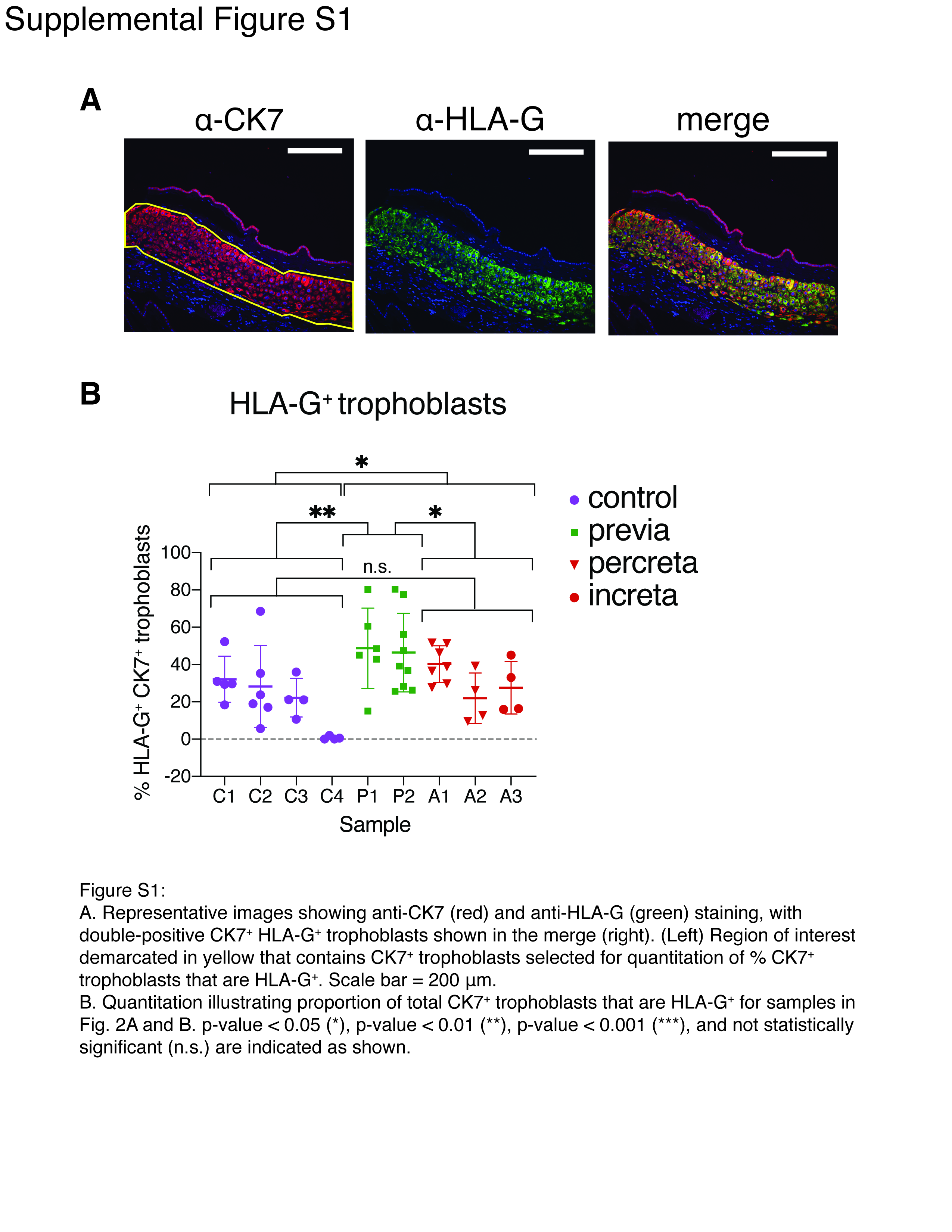

### Supplemental Fig. S2

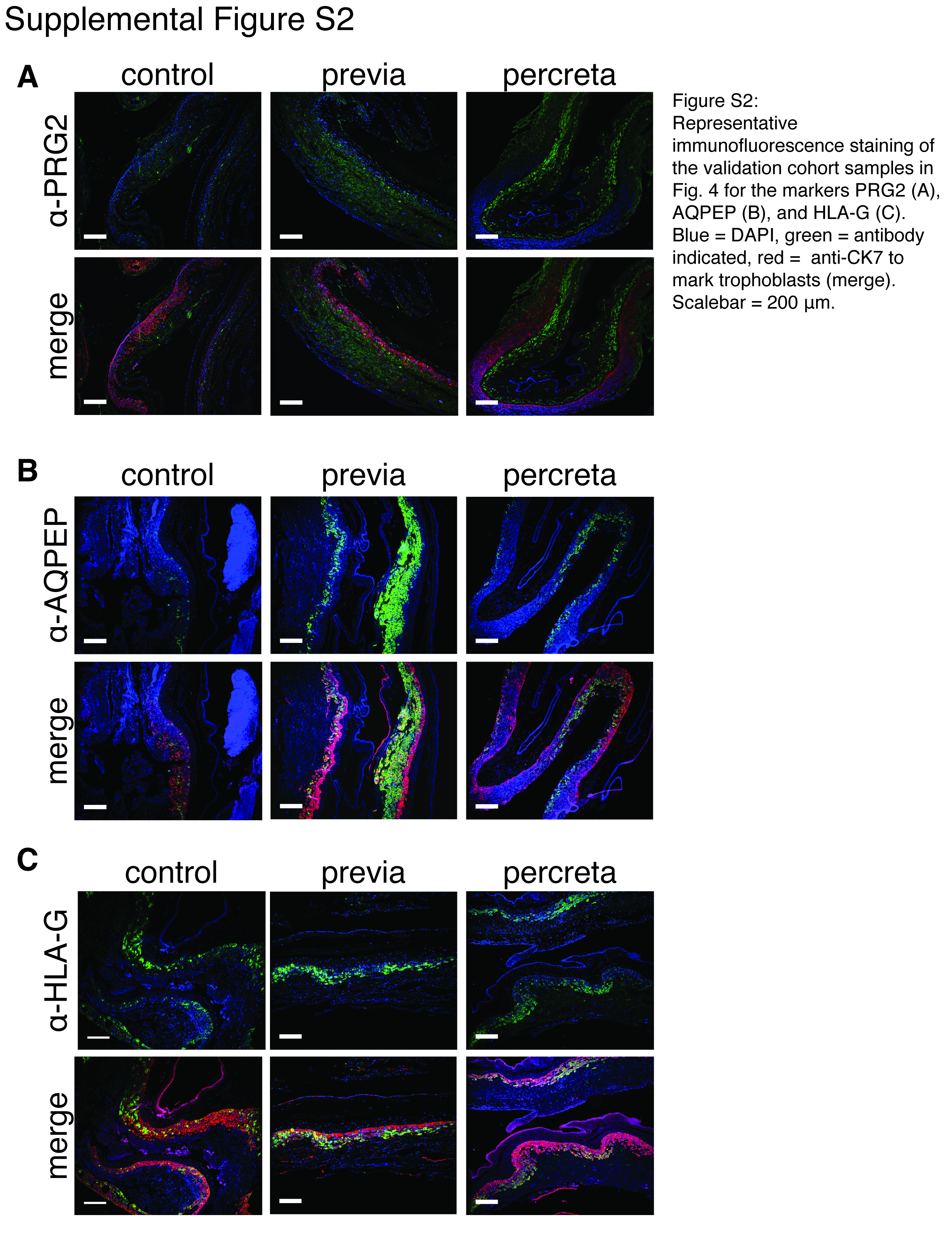
