## Supplemental Table S1 for "PRG2 and AQPEP are misexpressed in fetal membranes in placenta previa and percreta"

Cohort 1:

| Sample ID | Patient ID | Antepartum diagnosis | Pathology diagnosis | Previa, Y/N | Maternal age | GA at delivery (nearest half-week) | Placental weight | # prior uterine procedures (CS, D&C, and/or myomector | Mode of delivery | Other notes |
|---|---|---|---|---|---|---|---|---|---|---|
| C1 | NP1 | Spontaneous pre-term birth | Normal placenta | N | < 35 | 29.5 wk | 253 g | 1 | VD | |
| C2 | NP2 | Spontaneous pre-term birth | Placenta with accessory lobe | N | < 35 | 31 wk | 278 g, incl. accessory lobe | 0 | CD | advanced cervial dilation before C-section |
| C3 | NP3 | Spontaneous pre-term birth | Normal placenta | N | < 35 | 28 wk | 230 g | 0 | VD | vaginal bleeding; GBS (+) |
| C4 | NP4 | Spontaneous pre-term birth; posterior placenta | Normal placenta | N | ≥ 35 | 32 wk | 390 g (75th percentile) | 0 | VD | |
| C5 | NP5 | Spontaneous pre-term birth | Normal placenta; translucent fetal membranes with several fibrinous clots | N | < 35 | 31 wk | 460 g | 1 | CD | history of prior classical C-section |
| C6 | NP6 | Spontaneous pre-term birth | Large placenta for gestational age | N | < 35 | 28.5 wk | 312 g | 2 | CD | |
| C7 | NP7 | Spontaneous pre-term birth; probable placenta abruption | Placental disk with adjacent hemorrhage, consistent with placental abruption | N | < 35 | 29.5 wk | 330 g | 1 | VD | |
| C8 | NP8 | Spontaneous pre-term birth | Normal placenta | N | < 35 | 29 wk | 320 g | 1 | CD | |
| C9 | NP9 | Spontaneous pre-term birth | Amniotic fluid infection sequence; ruptured fetal membranes | N | < 35 | 28 wk | 278 g (75th-90th percentile) | 0 | VD | |
| P1 | NP10 | Marginal placenta previa | Placenta previa; premature villous maturation consistent with placental insufficiency and vascular disruption | Y | < 35 | 28 wk | 200 g | 0 | CD | vaginal bleeding |
| P2 | NP11 | Complete posterior placenta previa, bleeding | Placenta previa | Y | ≥ 35 | 29.5 wk | 247 g (10th-25th percentile) | 1 | CD | vaginal bleeding; high transverse C-section |
| A1 | NP12 | Complete anterior placenta previa; suspected percreta | Placenta percreta and previa; uterine rupture; fetal membranes adherent to uterine walls | Y | ≥ 35 | 30 wk | 200 g | 2 | CD | vaginal bleeding; gravid uterus with multiple omental and bladder adhesions |
| A2 | NP6 | Placenta previa and percreta | Placenta percreta and previa; uterine rupture | Y | < 35 | 30 wk | 207 g | ≥3 | CD | s/p Cesarean-hysterectomy |
| A3 | NP13 | Complete central placenta previa; concern for focal accreta; PPROM | Placenta increta and previa | Y | ≥ 35 | 30 wk | 220 g | ≥3 | CD | vaginal bleeding; s/p Cesarean hysterectomy |
| A4 | NP4 | Complete placenta previa with concern for accreta | Placenta percreta and previa | Y | < 35 | 32 wk | N/A | ≥3 | CD | s/p Cesarean hysterectomy |

Cohort 2:

| Sample ID | Patient ID | Antepartum diagnosis | Pathology diagnosis | Previa, Y/N | Maternal age | GA at delivery (nearest half-week) | Placental weight | # prior uterine procedures (CS, D&C, and/or myomector | Mode of delivery | Other notes |
|---|---|---|---|---|---|---|---|---|---|---|
| C10 | NP15 | Cervical cancer; preterm labor | Normal placenta; invasive, adenosquamous cancer | N | ≥ 35 | 35 wk | 400 g (50th-75th percentile) | 1 | CD | cervical cancer; s/p Cesarean-Hysterectomy |
| C11 | NP16 | Cervical cancer; scheduled Cesarean hysterectomy | Normal placenta; negative for residual carcinoma | N | < 35 | 34 wk | 345 g (25th-50th percentile) | 0 | CD | cervical cancer; s/p Cesaraean hysterectomy |
| C12 | NP17 | PPROM; Twin gestation | Small for gestataional age monochorionic diamniotic twin placenta | N | < 35 | 32.5 wk | 205.5 g | 0 | VD | GBS (+) |
| A6 | NP17 | PPROM; Twin gestation | Small for gestataional age monochorionic diamniotic twin placenta; microscopic placenta accreta | N | < 35 | 32.5 wk | 134 g | 0 | VD | GBS (+) |
| C13 | NP18 | PPROM | Intermediate acute chorioamnionitis | N | < 35 | 37 wk | 340 g (5th-10th percentile) | 0 | CD | CD due to arrest of dilation |
| C14 | NP19 | Di/Di twin pregnancy; Vaginal bleeding secondary to placenta previa with accreta, with concern for percreta; anterior placenta | Normal twin placenta | Y | ≥ 35 | 31.5 wk | Placenta B: 217 g (10-25th percentile) | 0 | CD | vaginal bleeding; IVF pregnancy; s/p Cesarean hysterectomy |
| A11 | NP19 | Di/Di twin pregnancy; Vaginal bleeding secondary to placenta previa with accreta, with concern for percreta; complete central placenta previa with concern for accreta | Placenta increta and previa; twin placenta | Y | ≥ 35 | 31.5 wk | N/A | 0 | CD | vaginal bleeding; IVF pregnancy; s/p Cesarean hysterectomy |
| P3 | NP20 | Placenta previa | Placenta previa | Y | ≥ 35 | 37 wk | 475 g (50th-75th percentile) | 1 | CD | |
| P4 | NP21 | Marginal placenta previa | Placenta previa; myometrium with implantation site changes | Y | < 35 | 36 wk | 500 g (50th-75th percentile) | 0 | CD | vaginal bleeding; s/p Cesarean hysterectomy |
| P5 | NP22 | Placenta previa with suspected accreta | Placenta previa; large placenta for gestational age | Y | ≥ 35 | 35 wk | 520.1 g (>97th percentile) | 0 | CD | |
| P6 | NP23 | Complete anterior placenta previa | Placenta previa; patchy chorioamnionic hemosiderosis suggestive of retroplacental bleeding | Y | ≥ 35 | 29.5 wk | 330 g (50th-75th percentile) | 3 | CD | |
| P7 | NP24 | Complete placenta previa | Placenta previa; large for gestational age placenta; disrupted maternal surface | Y | ≥ 35 | 34 wk | 500 g | 0 | CD | vaginal bleeding |
| P8 | NP25 | Complete placenta previa | Placenta previa | Y | <35 | 31 wk | 330 g (50th-75th percentile) | 2 | CD | vaginal bleeding |
| A5 | NP26 | Complete placenta previa and potential percreta | Placenta accreta (focal) and previa; laminar necrosis of decidua capsularis in fetal membranes | Y | ≥ 35 | 31 wk | N/A | ≥3 | CD | s/p Cesarean Hysterectomy |
| A7 | NP27 | Complete anterior placenta previa; placenta accreta | Placenta accreta (focal) and previa; chronic villitis | Y | < 35 | 33 wk | 400 g (75th-90th percentile) | 2 | CD | s/p Cesarean hysterectomy |
| A9 | NP28 | Placenta previa, possible accreta | Placenta accreta and previa | Y | ≥ 35 | 34 wk | N/A | 1 | CD | vaginal bleeding; s/p Cesarean hysterectomy |
| A10 | NP29 | Complete central placenta previa; no accreta | Placenta accreta and previa (cervical); small placenta for gestational age; laminar necrosis of decidua capsularis | Y | ≥ 35 | 30 wk | 197 g (5th-10th percentile) | 1 | CD | vaginal bleeding; s/p Cesarean hysterectomy |
| A12 | NP30 | Marginal anterior previa; placenta accreta | Placenta increta and previa; patchy non-specific chronic villitis; chronic deciduitis; amniotic fluid infection sequence | Y | ≥ 35 | 36 wk | 405 g (25th-50th percentile) | ≥3 | CD | mild preeclampsia; s/p Cesarean hysterectomy |
| A13 | NP31 | Placenta previa with suspected accreta, possibly percreta | Placenta percreta and previa | Y | < 35 | 34 wk | N/A | 2 | CD | |
| A14 | NP32 | Complete central placenta previa with suspicion for accreta or increta | Placenta percreta and previa (cervical) | Y | ≥ 35 | 36.5 wk | N/A | ≥3 | CD | s/p Cesarean hysterectomy |
| A15 | NP33 | Placenta previa with accreta, suspected percreta | Placenta percreta and previa; organizing intervillous hemorrhage; fetal membranes adherent to uterine cavity | Y | ≥ 35 | 34.5 wk | N/A | ≥3 | CD | s/p Cesarean Hysterectomy |
| A16 | NP34 | Placenta previa; placenta accreta with high suspicion of percreta | Placenta percreta and previa | Y | ≥ 35 | 35 wk | 396 g (25th-50th percentile) | ≥3 | CD | s/p Cesarean hysterectomy |
| A17 | NP35 | Placenta previa with concern for placenta accreta | Placenta percreta and previa | Y | ≥ 35 | 34 wk | N/A | ≥3 | CD | s/p Cesarean hysterectomy |
| A18 | NP36 | Placenta previa and percreta | Placenta percreta and previa (complete) | Y | ≥ 35 | 32 wk | N/A | 1 | CD | s/p Cesarean hysterectomy |
| A19 | NP37 | Complete central previa and accreta/ increta with suspected percreta | Placenta percreta and previa | Y | < 35 | 34 wk | N/A | ≥3 | CD | vaginal bleeding; s/p Cesarean hysterectomy |
| A20 | NP38 | Complete placenta previa and Increta | Placenta percreta; maternal placental underperfusion | Y | ≥ 35 | 36 wk | N/A | 1 | CD | |
| A21 | NP39 | Complete central placenta previa; placental placenta increta or percreta | Placenta percreta and previa | Y | < 35 | 28.5 wk | N/A | ≥3 | CD | Vaginal bleeding; s/p Cesarean hysterectomy |
| A22 | NP40 | Possible focal accreta | Placenta percreta | N | ≥ 35 | 37 wk | N/A | 2 | CD | |

Additional patient details available upon request.
