## Supplemental Table S2 for "PRG2 and AQPEP are misexpressed in fetal membranes in placenta previa and percreta"

| Sample number | Patient ID | Gestational Age (weeks at birth) | Diagnosis | # of reads | % aligned | # of uniquely-mapped reads |
|---|---|---|---|---|---|---|
| C1 | NP1 | 28-32 | SPTB | 27142401 | 62.1 | 11663958 |
| C2 | NP2 | 28-32 | SPTB | 19083626 | 84.3 | 9668586 |
| C3 | NP3 | 28-32 | SPTB | 22233745 | 65.3 | 7912656 |
| C4 | NP4 | 28-32 | SPTB | 17997868 | 85.7 | 9717229 |
| C5 | NP5 | 28-32 | SPTB | 21174067 | 66.2 | 9251373 |
| C6 | NP6 | 28-32 | SPTB | 24277580 | 81.4 | 11797884 |
| C7 | NP7 | 28-32 | SPTB | 28120419 | 73.2 | 13235606 |
| C8 | NP8 | 28-32 | SPTB | 27775683 | 68.1 | 12389482 |
| C9 | NP9 | 28-32 | SPTB | 27861742 | 77.2 | 14497244 |
| P1 | NP10 | 28-32 | Previa | 24415440 | 79.7 | 12570582 |
| P2 | NP11 | 28-32 | Previa | 18513937 | 77.2 | 9476099 |
| A1 | NP12 | 28-32 | Percreta/Previa | 23014613 | 81.3 | 10927154 |
| A2 | NP6 | 28-32 | Percreta/Previa | 24647905 | 81.7 | 11216497 |
| | | | Average: | 23558387 | 75.6 | 11101873 |
