## Supplemental Table S3 for "PRG2 and AQPEP are misexpressed in fetal membranes in placenta previa and percreta"

| Official gene symbol | Gene name | log2 fold change | p-value | p-adjusted | log2 EVT / VCT (Vondra et al. 2019) | Cell type (Tsang et al. 2017) | log2 signal EVT - VCT (Apps et al. 2009) |
|---|---|---|---|---|---|---|---|
| ACCN1 | Amiloride-Sensitive Cation Channel 1, Neuronal | 5.453668238 | 2.62E-08 | 3.27E-05 | N/A | N/A | N/A |
| LOC284379 | solute carrier family 7 member 3 pseudogene | 5.075388315 | 1.17E-06 | 0.00061 | N/A | N/A | N/A |
| AADACL3 | arylacetamide deacetylase like 3 | 3.938506191 | 1.84E-08 | 3.16E-05 | N/A | N/A | N/A |
| AQPEP | Aminopeptidase Q (Laeverin) | 3.536252728 | 6.23E-10 | 5.50E-06 | 2.70233 | Extravillous trophoblast | 5.08 |
| MYL4 | myosin light chain 4 | 3.518454207 | 1.23E-05 | 0.003517 | N/A | N/A | N/A |
| LY6D | lymphocyte antigen 6 complex, locus D | 3.461477371 | 9.15E-08 | 9.11E-05 | N/A | Extravillous trophoblast | N/A |
| PRG2 | proteoglycan 2, pro eosinophil major basic protein | 3.286498673 | 4.97E-07 | 0.000419 | 2.77165 | N/A | 3.902415745 |
| PIPOX | pipecolic acid and sarcosine oxidase | 3.224391314 | 2.69E-08 | 3.27E-05 | 2.35136 | N/A | 3.576006688 |
| NXPH3 | neurexophilin 3 | 3.030988073 | 1.07E-06 | 0.000583 | N/A | N/A | N/A |
| HUNK | hormonally up-regulated Neu-associated kinase | 2.941975801 | 2.58E-09 | 5.65E-06 | 1.24193 | N/A | N/A |
| SLC30A2 | solute carrier family 30 member 2 | 2.813119832 | 1.51E-09 | 5.50E-06 | N/A | N/A | -3.509387331 |
| PHGDH | phosphoglycerate dehydrogenase | 2.76008125 | 8.49E-08 | 9.11E-05 | N/A | N/A | N/A |
| HLA-G | major histocompatibility complex, class I, G | 2.7089126 | 9.94E-06 | 0.003199 | 3.33738 | Extravillous trophoblast | 6.088389436 |
| SLC6A2 | solute carrier family 6 member 2 | 2.697195881 | 3.22E-06 | 0.001381 | 2.48436 | Extravillous trophoblast | N/A |
| ISM2 | isthmin 2 | 2.652772225 | 0.000127 | 0.022438 | 2.01627 | N/A | N/A |
| NES | nestin | 2.634552179 | 2.02E-08 | 3.16E-05 | N/A | N/A | N/A |
| AGTR1 | angiotensin II receptor type 1 | 2.595282999 | 3.28E-06 | 0.001381 | N/A | N/A | N/A |
| HTRA4 | HtrA serine peptidase 4 | 2.554869856 | 7.67E-07 | 0.000544 | 3.16153 | N/A | 4.381783065 |
| FHOD3 | formin homology 2 domain containing 3 | 2.550517482 | 9.10E-05 | 0.017472 | 1.92231 | N/A | N/A |
| KIF12 | kinesin family member 12 | 2.544470589 | 3.56E-06 | 0.001444 | N/A | N/A | N/A |
| CILP2 | cartilage intermediate layer protein 2 | 2.541152787 | 0.000107 | 0.019506 | N/A | N/A | N/A |
| REPS2 | RALBP1 associated Eps domain containing 2 | 2.538520387 | 2.20E-07 | 0.0002 | 3.22794 | N/A | 2.727446925 |
| PYCR1 | pyrroline-5-carboxylate reductase 1 | 2.535821071 | 1.35E-05 | 0.00359 | 3.46776 | Extravillous trophoblast | 2.510180705 |
| NOTUM | NOTUM, palmitoleoyl-protein carboxylesterase | 2.535664567 | 1.19E-05 | 0.003517 | 3.0203 | Extravillous trophoblast | 5.006430495 |
| C2orf55 | Uncharacterized Protein KIAA1211-Like | 2.528428804 | 3.95E-05 | 0.009617 | N/A | N/A | N/A |
| AIF1L | allograft inflammatory factor 1 like | 2.394679642 | 1.24E-05 | 0.003517 | 3.19369 | Extravillous trophoblast | N/A |
| LIFR | leukemia inhibitory factor receptor alpha | 2.393404838 | 2.58E-09 | 5.65E-06 | 1.04582 | N/A | N/A |
| DIO2 | deiodinase, iodothyronine type II | 2.321422422 | 6.07E-05 | 0.014144 | N/A | N/A | 3.780584281 |
| LIPG | lipase G, endothelial type | 2.293775855 | 0.00015 | 0.025177 | 1.51376 | N/A | N/A |
| VWCE | von Willebrand factor C and EGF domains | 2.286392884 | 8.87E-06 | 0.003035 | N/A | N/A | N/A |
| ANKRD58 | Ankyrin Repeat Domain-Containing Protein SOWAHD | 2.174652501 | 8.53E-05 | 0.016667 | N/A | N/A | N/A |
| ITM2B | integral membrane protein 2B | 2.164683089 | 1.26E-06 | 0.000629 | 2.36328 | N/A | N/A |
| CSF2RB | colony stimulating factor 2 receptor beta common subunit | 2.13140814 | 1.87E-05 | 0.004861 | 2.55475 | N/A | N/A |
| HES2 | hes family bHLH transcription factor 2 | 2.08094067 | 2.11E-05 | 0.00537 | N/A | N/A | N/A |
| ADAM19 | ADAM metallopeptidase domain 19 | 2.05370837 | 8.25E-07 | 0.000544 | 3.07399 | N/A | 5.997506501 |
| BCAR4 | breast cancer anti-estrogen resistance 4 | 2.041573601 | 0.000205 | 0.032094 | 2.51665 | Extravillous trophoblast | N/A |
| ANKRD24 | ankyrin repeat domain 24 | 2.039778058 | 1.57E-06 | 0.000747 | N/A | N/A | N/A |
| HEXB | hexosaminidase subunit beta | 2.034473708 | 1.31E-09 | 5.50E-06 | N/A | N/A | N/A |
| LYVE1 | lymphatic vessel endothelial hyaluronan receptor 1 | 2.027426026 | 2.26E-05 | 0.005618 | N/A | Hofbauer cell | N/A |
| FLT1 | fms related tyrosine kinase 1 | 2.027299198 | 5.05E-06 | 0.001843 | 2.01911 | N/A | N/A |
| MAG | myelin associated glycoprotein | 1.934656289 | 0.000119 | 0.021409 | N/A | N/A | 2.417970761 |
| PTN | pleiotrophin | 1.917978915 | 0.000181 | 0.029161 | 1.79807 | N/A | N/A |
| CPM | carboxypeptidase M | 1.911118809 | 8.65E-07 | 0.000544 | 1.85138 | N/A | N/A |
| FLJ23867 | uncharacterized protein FLJ23867 | 1.89893325 | 0.000215 | 0.033171 | N/A | N/A | N/A |
| AIFM3 | apoptosis inducing factor, mitochondria associated 3 | 1.886609305 | 4.63E-05 | 0.01102 | N/A | N/A | N/A |
| JAM2 | junctional adhesion molecule 2 | 1.872464334 | 9.44E-07 | 0.000544 | 2.82357 | N/A | 4.019202123 |
| DHCR24 | 24-dehydrocholesterol reductase | 1.840885874 | 1.25E-05 | 0.003517 | -1.29498 | N/A | N/A |
| LPAR2 | lysophosphatidic acid receptor 2 | 1.823294381 | 6.93E-05 | 0.015346 | N/A | N/A | N/A |
| HN1 | hematological and neurological expressed 1 | 1.8191031 | 3.01E-06 | 0.00137 | 2.22452 | N/A | N/A |
| GPR146 | G protein-coupled receptor 146 | 1.740346686 | 9.30E-07 | 0.000544 | 2.52166 | N/A | N/A |
| MCAM | melanoma cell adhesion molecule | 1.726067275 | 6.51E-07 | 0.000509 | 2.58827 | N/A | 3.051436173 |
| BGN | biglycan | 1.71396756 | 0.000145 | 0.024774 | 2.22116 | N/A | N/A |
| HTRA1 | HtrA serine peptidase 1 | 1.709591331 | 1.15E-05 | 0.003517 | 2.89865 | N/A | 2.680932036 |
| IL2RB | interleukin 2 receptor subunit beta | 1.697849443 | 0.000349 | 0.049626 | 2.97153 | N/A | 5.299850104 |
| ITM2C | integral membrane protein 2C | 1.665494141 | 7.01E-05 | 0.015346 | N/A | N/A | N/A |
| TLE6 | transducin like enhancer of split 6 | 1.570280057 | 9.23E-06 | 0.003062 | N/A | N/A | N/A |
| ENDOG | endonuclease G | 1.543312366 | 5.02E-06 | 0.001843 | N/A | N/A | N/A |
| CCNYL1 | cyclin Y like 1 | 1.528492739 | 8.40E-05 | 0.016667 | 1.03977 | N/A | N/A |
| HSPG2 | heparan sulfate proteoglycan 2 | 1.511961776 | 1.34E-05 | 0.00359 | 2.33221 | N/A | N/A |
| PRRG4 | proline rich and Gla domain 4 | 1.502809482 | 9.27E-05 | 0.017487 | N/A | N/A | N/A |
| MGAT5 | mannosyl | 1.493887051 | 0.000221 | 0.033604 | 2.83196 | N/A | N/A |
| ACVRL1 | activin A receptor like type 1 | 1.491887642 | 0.000102 | 0.018954 | N/A | N/A | 5.009335024 |
| EPHB2 | EPH receptor B2 | 1.488126387 | 5.86E-06 | 0.00207 | N/A | N/A | 2.444690526 |
| TPM1 | tropomyosin 1 | 1.453669428 | 4.67E-06 | 0.001823 | 2.43131 | N/A | 2.631045926 |
| POMC | proopiomelanocortin | 1.433317937 | 0.0003 | 0.043788 | N/A | N/A | N/A |
| FGFR1 | fibroblast growth factor receptor 1 | 1.404246179 | 8.36E-05 | 0.016667 | 1.73415 | N/A | N/A |
| KIF21A | kinesin family member 21A | 1.373069931 | 0.000174 | 0.028446 | 1.93513 | N/A | N/A |
| UBE2A | ubiquitin conjugating enzyme E2 A | 1.346751546 | 7.19E-05 | 0.015438 | N/A | N/A | N/A |
| QSOX1 | quiescin sulfhydryl oxidase 1 | 1.329113869 | 6.56E-05 | 0.014951 | 2.81357 | N/A | 3.191120421 |
| ICK | intestinal cell kinase | 1.320238452 | 7.77E-05 | 0.016036 | N/A | N/A | N/A |
| EFNB1 | ephrin B1 | 1.247550708 | 0.000171 | 0.02837 | 2.44074 | N/A | 2.4603697 |
| SOX4 | SRY-box 4 | 1.005782298 | 0.000226 | 0.033888 | 1.57694 | N/A | N/A |
| SH3PXD2A | SH3 and PX domains 2A | 1.003157181 | 7.55E-05 | 0.015881 | 1.7425 | N/A | N/A |
| ZNF704 | zinc finger protein 704 | 0.954418118 | 0.000185 | 0.029404 | 1.53469 | N/A | N/A |
| RNF19B | ring finger protein 19B | -1.161219723 | 0.00013 | 0.022662 | -0.758406 | N/A | N/A |
| NIPA1 | non imprinted in Prader-Willi/Angelman syndrome 1 | -2.904374574 | 0.000272 | 0.040193 | N/A | N/A | N/A |
| ADAMDEC1 | ADAM like decysin 1 | -2.948596404 | 0.000305 | 0.043931 | N/A | N/A | N/A |
| | | | | Vondra et al. EVT max | 4.81 | Apps et al. highest value | 6.67 |
| | | | | Vondra et al. VCT max | -5.42 | Apps et al. lowest value | -7.06 |
